# Season-long head acceleration exposure, prior contact-sport history, and cognitive performance in British amateur American football players

**DOI:** 10.64898/2026.09.30.755702

**Authors:** Paul Weightman Potter, Emily Rose, Nicholas Tolliday, Bert Bond

**Affiliations:** Faculty of Health and Life Sciences, Department of Clinical and Biomedical Sciences, University of Exeter Medical School, St Luke’s Campus, Exeter, EX1 2LU; Exeter Head Impacts, Brain Injury and Trauma (ExHIBIT) research group

**Keywords:** American football, head acceleration, cognitive performance, repeated assessment, observational study

## Abstract

**Background:** Head acceleration events (HAEs) are common in contact sports, but season-long exposure and its relationship with cognitive performance in amateur British American football remain poorly characterised.

**Methods:** Thirty male players were recruited from three British amateur American football teams. Selected Sport Concussion Assessment Tool 6 (SCAT6) components and computerised tasks were assessed across the season. Seasonal analyses included 20 participants, and exposure-linked analyses included 17. Helmet-sensor events were attributed to participants and restricted to participant-assignment and baseline-to-post-season calendar windows.

Mixed-effects models assessed seasonal outcomes. Principal component analysis (PCA) summarised standardised baseline-to-post-season changes, and six exposure associations were evaluated.

**Results:** Players recorded a mean of 768 (SD 482) events. Several individual tests improved across the season. The first principal component (PC1) explained 21.9% of change variance and had loadings with opposing signs. Prior seasons had a nominal association with PC1 (slope −0.153, 95% CI −0.305 to −0.00018; r^2^=0.233; p=0.0498), which did not survive six-test Benjamini–Hochberg (BH) adjustment (p=0.299). No in-season exposure association survived adjustment.

**Conclusion:** Repeated-test performance improved, while exposure associations remained uncertain. The mixed PC1 pattern cannot be interpreted as a uniform measure of global cognitive worsening or improvement.

## Introduction

Head acceleration events (HAEs) are frequent in contact sports such as American football, rugby, and combat sports. Collegiate American football players can experience hundreds of recorded events per season [1]. Head motion involves linear and rotational kinematics, and sufficiently severe loading can contribute to neural and vascular injury [2,3]. Traumatic brain injury carries a substantial public-health burden [4,5]. Field measurements of exposure therefore complement clinical assessment.

Many HAEs do not result in overt clinical symptoms, which makes exposure difficult to track through symptom reporting alone. Nevertheless, repetitive head impacts accumulated over the course of a playing career have been implicated in longer-term neurological complications, including chronic traumatic encephalopathy (CTE) [6]. Recognition of symptomatic injury is further complicated by athletes underreporting symptoms due to fears of missing playing time [7]. Recent advances in wearable technology have enabled studies that quantify HAEs in other contact sports such as rugby [8–10].

The short-to medium-term cognitive effects of repetitive head impacts remain incompletely characterised. Collegiate and high-school American football studies have reported changes in memory, executive function, and neurophysiological measures across a season, including among athletes without diagnosed concussion [11–13]. Repeated testing and individual characteristics complicate interpretation of cognitive performance [14,15]. A controlled amateur boxing study also reported altered cerebral autoregulation after boxing [16], and imaging studies have identified season-level changes in white-matter and functional measures [17,18]. These findings support combining exposure monitoring with clearly defined functional outcomes.

British American football provides an important but under-studied context for HAE research. The sport is played at youth, university, and amateur adult levels, yet UK-specific data describing the frequency and magnitude of HAEs across a season and their relationship with cognitive performance are limited. Compared with American collegiate or professional settings, British teams often have shorter seasons and less frequent training, which may reduce exposure. However, amateur teams may also have less access to medical, monitoring, and performance-support infrastructure. Characterising HAE exposure and cognitive performance in this setting is therefore important for player welfare and for informing future monitoring strategies.

This study aimed to characterise season-long recorded helmet-sensor exposure in British amateur American football and examine its relationship with cognitive change. A secondary aim was to examine prior contact-sport history, recorded as previous seasons played. We hypothesised that greater in-season exposure and greater prior contact-sport history would be associated with less favourable cognitive outcomes. The analyses are observational and exploratory.

## Methods

### Participants

Thirty adult male players were recruited from three British amateur American football teams, comprising two university teams and one adult team. Nine participants withdrew and were excluded entirely. One of the remaining participants had exposure data only, leaving 20 for seasonal cognitive analyses, with outcome-specific missingness. Three of these 20 were excluded from exposure-linked analyses because sensor recording was incomplete, leaving 17 linked participants. Baseline age in the 20-person cognitive cohort was 22.8 (SD 3.8) years, range 19–32. Sensor exclusions were distinguished from withdrawal and individual missing measurements. All participants provided written informed consent. University of Exeter ethics approval was granted on 1 September 2024 (reference 6856378). Exclusion criteria were a diagnosed concussion within three months before baseline, neurological disease, diabetes, cardiovascular disease, renal disease, or hepatic disease.

### Study design

This was a season-long observational study with baseline, mid-season, and post-season assessments. Scheduling reflected the different university and adult-team seasons; mid-season assessments therefore occurred at different stages of the teams’ calendars. All post-season measures occurred within two weeks of each participant’s last match. One participant completed baseline and post-season assessments only; a separate assessment outside this study was excluded. Participant and session assignments were verified against the participant key, investigator records, and recording timestamps. Seventeen participants had a study mid-season assessment.

### Head acceleration event measurement

Helmet-mounted tri-axial accelerometers (ACT Tracker, Northern Sports Insight and Intelligence Oy, Finland) recorded linear acceleration during contact training and matches [19]. The sensor trigger was acceleration above 10 g on an axis, sampling was 1 kHz, and the nominal event window was 75 ms. Exported resultant acceleration traces (labelled Total) contained 76 samples labelled 0–75 ms. Event Gadd Severity Index (GSI) was the sum of all 76 acceleration-in-g samples raised to the power 2.5, multiplied by 0.001 s. Participant GSI summed retained event values. GSI was calculated using this rectangular-sum approximation.

Events were grouped by peak recorded acceleration into 10 to <40 g, 40 to <80 g, and ≥80 g; exactly 80 g entered the highest category. The total number of HAEs was also recorded. These are recorded helmet-sensor events, not independently verified head impacts.

Players activated and deactivated their sensors. Processing preserved exact profile/sensor labels and documented transfers, removed exact duplicate records, and restricted events to verified participant-assignment and baseline/post-season calendar windows. No event-level artefact threshold or screening was applied. Sensor-recording sufficiency was assessed individually, and three participants were excluded from exposure-linked analyses due to self-reported incomplete data collection.

Prior contact-sport history was the self-reported number of previous seasons across contact sports. Prior seasons represent sporting history rather than a directly measured lifetime impact dose.

### Cognitive testing

Assessments used selected SCAT6 components and PsyToolkit tasks [20–27]. The assessed components were symptoms, orientation, immediate memory, delayed recall, digits backward, reverse months/concentration, and modified Balance Error Scoring System (mBESS) errors. Tandem gait was not administered; this was not the full SCAT6. Delayed recall occurred at least five minutes after immediate recall. Task presentation timings were not recorded. Raw questionnaire responses were decoded with instrument-specific rules.

Symptom count and severity used 22 ratings from 0 to 6. Summary scores were derived from complete, valid component responses. Immediate memory counted words across three trials; concentration combined digits backward and reverse months; mBESS summed three stance-error counts. Missing or invalid responses were not converted to zero.

### Statistical analysis

Seasonal outcomes were analysed in R [28] using available observations without imputation. Time was included as a categorical fixed effect, with a participant-specific random intercept. Models were fitted using lmerTest with restricted maximum likelihood (REML) and assumed compound symmetry, with Satterthwaite Type III tests. Pairwise visit contrasts were estimated using emmeans with Satterthwaite degrees of freedom; a custom step-down Holm–Šídák function adjusted p values across three within-outcome visit comparisons.

Confidence intervals are marginal 95% intervals; adjustments to p values are specified separately. Descriptions report mean (SD), median (Q1–Q3), and analysis-specific denominators. Orientation had near-boundary participant variance: its time effect concerns mid-season departures because baseline and post-season scores were constant at 5.

### Principal component summary of change

For verified measures, raw change was post-season minus baseline. Change scores were oriented so that favourable change was positive: memory, orientation, digits backward, and concentration used post-season minus baseline; symptoms, errors, balance errors, and reaction times (RTs) used baseline minus post-season. The 17 change variables were standardised using observed means and sample SDs in the linked cohort. Constant/all-missing measures were removed. The primary task-scoring approach was accompanied by a correct-response RT sensitivity analysis. Stroop used mean incongruent minus congruent RT without an accuracy or block-label filter; the sensitivity used correct responses. Go/No-Go errors were summed, and the primary RT definition excluded exactly 2,000 ms; the sensitivity used correct Go responses only. Every 2-back file contained 75 trials; errors were calculated as 75 minus summed correctness, and RT was calculated from correct trials, excluding those of exactly 3,000 ms. Visual search used correct target-present trials identified by the prespecified task flag within each array size. No additional trimming, timeout rule, or practice exclusion was introduced.

PCA used 17 linked participants. Constant orientation change was removed, leaving 16 variables; one missing baseline-to-post-season 2-back RT change was mean-imputed for full-cohort PCA. Native PC1 scores and loadings were retained without whole-component sign alignment. Digits backward and concentration had identical changes and were both included in the primary variable set; a domain-balanced sensitivity averaged standardised changes within eight domains and then averaged the domain scores with equal weight, excluding separate digits-backward weighting and orientation (Supplementary Table S6). Loadings with opposing signs were inspected because aligning input directions does not necessarily produce a global cognition scale. PC1 is an exploratory pattern of cognitive, symptom, and balance changes [29].

### Exposure associations with the principal component

Six separate simple linear regressions used PC1 as outcome and, individually, prior seasons, total events, total GSI, and event counts at 10 to <40 g, 40 to <80 g, or ≥80 g as predictors. All used 17 complete exposure records. Slopes, marginal 95% CIs, r^2^, raw p values, and Benjamini–Hochberg (BH) p values across six tests are reported. Correct-response RT and domain-balanced analyses are sensitivities. Analytical choices were specified consistently across the reported outcomes and sensitivity analyses. Seasonal omnibus BH adjustment covered 17 outcomes separately for each scoring variant.

### Exploratory prediction

As a secondary exploratory analysis, nested cross-validation of penalised linear regression models fitted using glmnet [30,31] was used to assess whether the six exposure measures predicted the fixed full-cohort PC1. A sensitivity analysis re-estimated outcome processing and PCA within each training split and omitted total events because it equals the sum of the three event-category counts. Given the small sample, these analyses were treated as descriptive assessments of out-of-sample performance. Cross-validation procedures and performance metrics are reported in the Supplementary Material.

## Results

### Head acceleration event exposure over the course of a season

Thirty players were recruited, 20 contributed seasonal cognitive observations, and 17 contributed linked exposure/cognition data. The exposure-only participant was not included in the exposure summaries reported here, which describe the linked cohort. In the linked 17, mean exposure was 767.9 (SD 481.5) recorded events: 674.0 (439.6) at 10 to <40 g, 69.0 (38.9) at 40 to <80 g, and 24.9 (15.1) at ≥80 g. Mean total GSI was 20,199 (12,034) g^2^·^5^ s. Table 1 reports characteristics and exposure distributions.

**Table 1.** Characteristics and recorded exposure of the 17 linked participants. Continuous values are mean (SD), median (Q1–Q3), and range; categorical values are n (%). Q1 and Q3 are the first and third quartiles. Prior history is reported in seasons. GSI units are g^2^·^5^ s. Counts represent recorded helmet-sensor events, not independently verified impacts.

| Characteristic | n | Mean (SD) or n (%) | Median (Q1–Q3) | Range |
| --- | --- | --- | --- | --- |
| Age (years) | 17 | 22.4 (3.6) | 21.0 (20.0–25.0) | 19.0–32.0 |
| Prior contact-sport seasons | 17 | 10.6 (5.9) | 11.0 (9.0–15.0) | 1.0–21.0 |
| Total recorded events | 17 | 767.9 (481.5) | 618.0 (470.0–740.0) | 384.0–2,043.0 |
| Total GSI | 17 | 20,199 (12,034) | 18,110 (13,744–23,798) | 5,125–54,778 |
| Events 10 to <40 g | 17 | 674.0 (439.6) | 542.0 (398.0–616.0) | 328.0–1,855.0 |
| Events 40 to <80 g | 17 | 69.0 (38.9) | 60.0 (51.0–78.0) | 14.0–160.0 |
| Events ≥80 g | 17 | 24.9 (15.1) | 26.0 (15.0–29.0) | 4.0–65.0 |
| Male sex | 17 | 17 (100%) | — | — |

**Table 2.** Primary-scoring individual outcomes. Values are median (Q1–Q3), where Q1 and Q3 are the first and third quartiles. Most outcomes have n=20, 17, and 20; 2-back RT has n=19, 17, and 19. Missing observations were not imputed. Model estimates, marginal confidence intervals, and multiplicity adjustments are separate from these descriptive summaries.

| Outcome | Baseline n=20 | Mid-season n=17 | Post-season n=20 |
| --- | --- | --- | --- |
| Symptom count | 2.5 (0.8–5.2) | 2.0 (1.0–3.0) | 1.5 (0.8–3.0) |
| Symptom severity | 4.0 (0.8–6.5) | 2.0 (1.0–6.0) | 2.0 (0.8–3.0) |
| Orientation | 5.0 (5.0–5.0) | 5.0 (5.0–5.0) | 5.0 (5.0–5.0) |
| Immediate memory | 21.0 (20.0–23.2) | 24.0 (21.0–25.0) | 24.0 (20.0–25.0) |
| Delayed recall | 7.0 (6.0–8.2) | 8.0 (6.0–9.0) | 9.0 (8.0–9.2) |
| Digits backward | 2.0 (1.0–3.0) | 2.0 (1.0–3.0) | 3.0 (2.0–3.2) |
| Concentration | 3.0 (2.0–4.0) | 3.0 (2.0–4.0) | 4.0 (3.0–4.2) |
| mBESS errors | 3.0 (2.0–4.5) | 4.0 (1.0–6.0) | 3.5 (1.0–6.2) |
| Stroop interference (ms) | 66.1 (29.2–125.7) | 75.4 (–10.9–120.8) | 87.2 (34.4–159.1) |
| Go/No-Go errors | 0.0 (0.0–0.0) | 0.0 (0.0–1.0) | 0.0 (0.0–1.0) |
| Go/No-Go RT (ms) | 353.2 (326.3–415.4) | 344.1 (307.0–399.3) | 327.7 (305.0–344.9) |
| 2-back errors | 5.0 (2.8–9.2) | 1.0 (1.0–7.0) | 1.5 (1.0–4.0) |
| 2-back RT (ms) | 499.5 (472.0–682.5); n=19 | 491.9 (454.9–581.1); n=17 | 480.1 (409.4–545.8); n=19 |
| Search RT 5 items (ms) | 681.4 (603.3–741.6) | 646.0 (572.6–708.0) | 609.1 (526.8–654.4) |
| Search RT 10 items (ms) | 777.1 (709.9–907.1) | 731.2 (663.9–893.2) | 723.9 (618.8–771.9) |
| Search RT 15 items (ms) | 983.1 (917.5–1,209.2) | 879.3 (801.0–1,050.7) | 890.6 (757.5–933.4) |
| Search RT 20 items (ms) | 1,100.3 (954.2–1,243.4) | 884.6 (765.8–1,154.2) | 844.4 (757.8–1,024.5) |

### Cognitive test performance improved over the course of a season

Most outcomes had 20 baseline, 17 mid-season, and 20 post-season observations; 2-back RT had 19, 17, and 19. Available-observation models estimated baseline-to-post-season improvements in immediate memory (+2.35 words over three trials, 95% CI 0.84 to 3.86), delayed recall (+1.15, 0.57 to 1.73), 2-back errors (−3.10, −5.13 to −1.07), 2-back RT (−115.3 ms, −183.1 to −47.5), and 15- and 20-item search RT (−182.1 and −272.1 ms). These outcomes had both BH-adjusted omnibus and Holm–Šídák-adjusted baseline-to-post-season p<0.05. The 10-item search also met both criteria. The five-item search had an adjusted baseline-to-post-season p=0.0268 but 17-outcome omnibus BH p=0.0582. Concentration and digits had nominal time effects that did not survive omnibus adjustment. Primary-scoring omnibus tests and baseline-to-post-season contrasts, including the correct-response RT sensitivity contrasts, are reported in Tables S2, S3, and S3b.

PC1 explained 21.92% of change variance. Largest absolute weights were symptom severity (−0.442), symptom count (−0.433), Go/No-Go RT (−0.413), and 2-back errors (−0.408). Delayed recall and 10-item search had positive weights (+0.211 and +0.216). Favourable changes in different measures could therefore move PC1 in opposing directions. Correct-response RT produced a similar mixed pattern. This component does not substantiate uniform global cognitive worsening or improvement.

### Exposure associations with PC1

Prior seasons were nominally associated with PC1 (slope −0.1527 per season, 95% CI −0.3053 to −0.00018; r^2^=0.2329; raw p=0.0498), but six-test BH p=0.2986. None of the five in-season associations survived adjustment (Figure 2; Table S4). Total events had r^2^=0.0621, raw p=0.3347, and BH p=0.3984; GSI had r^2^=0.1487, raw p=0.1264, and BH p=0.3792. Bin-count r^2^ values ranged from 0.0480 to 0.1062, with BH p=0.3984. Negative slopes concern the native component axis, not evidence of protection. Correct-response RT and domain-balanced sensitivities also had no six-test BH p<0.05 (Tables S4 and S6).

**Figure 1.**
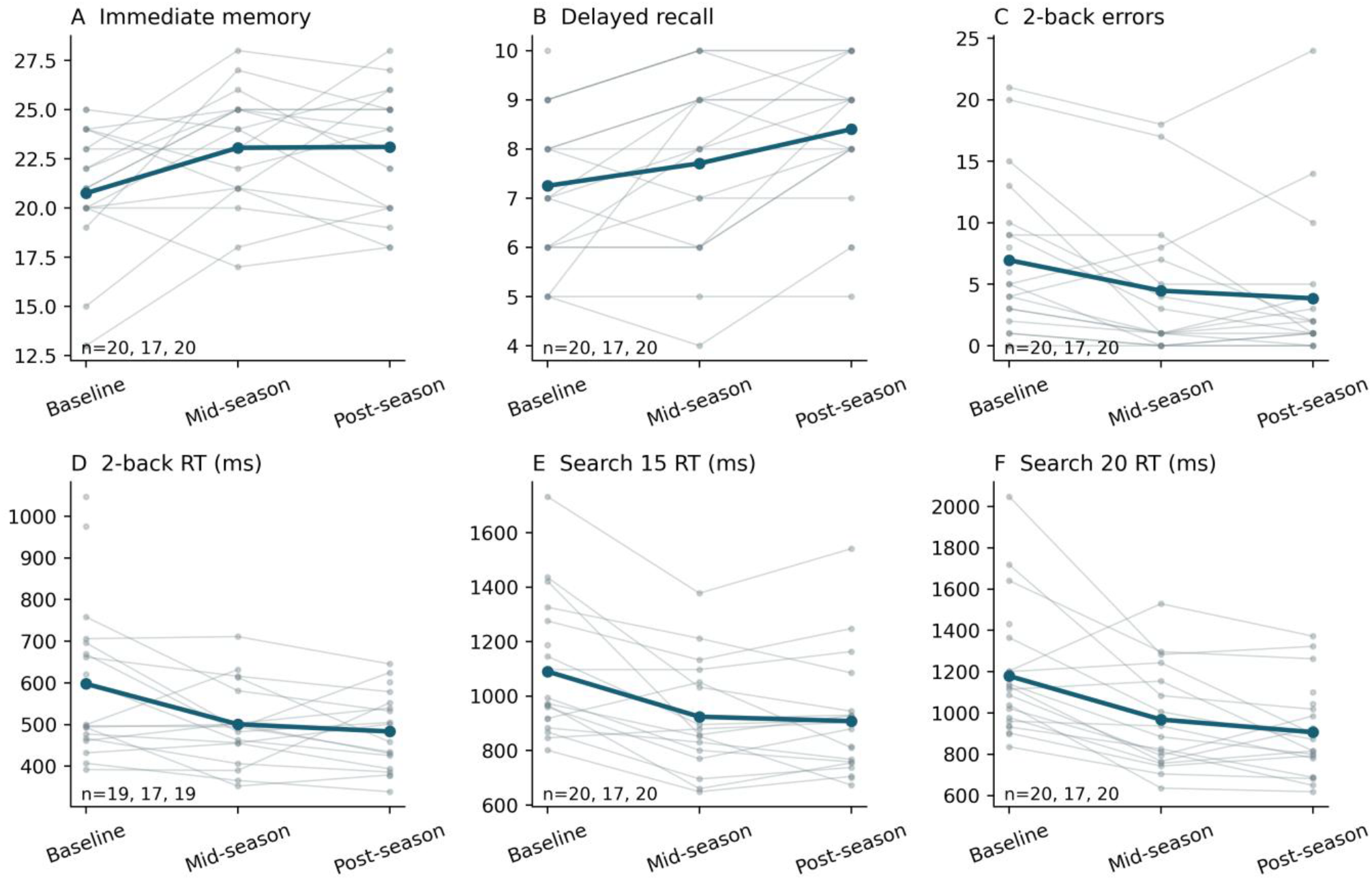
Repeated cognitive assessments in the 20-person cognitive cohort. Thin lines and points show observed participant trajectories; thick lines show observed means. Outcome-specific sample sizes are shown. Plots are descriptive; inference uses REML, Satterthwaite tests, and the multiplicity adjustments stated in the methods.

**Figure 2.**
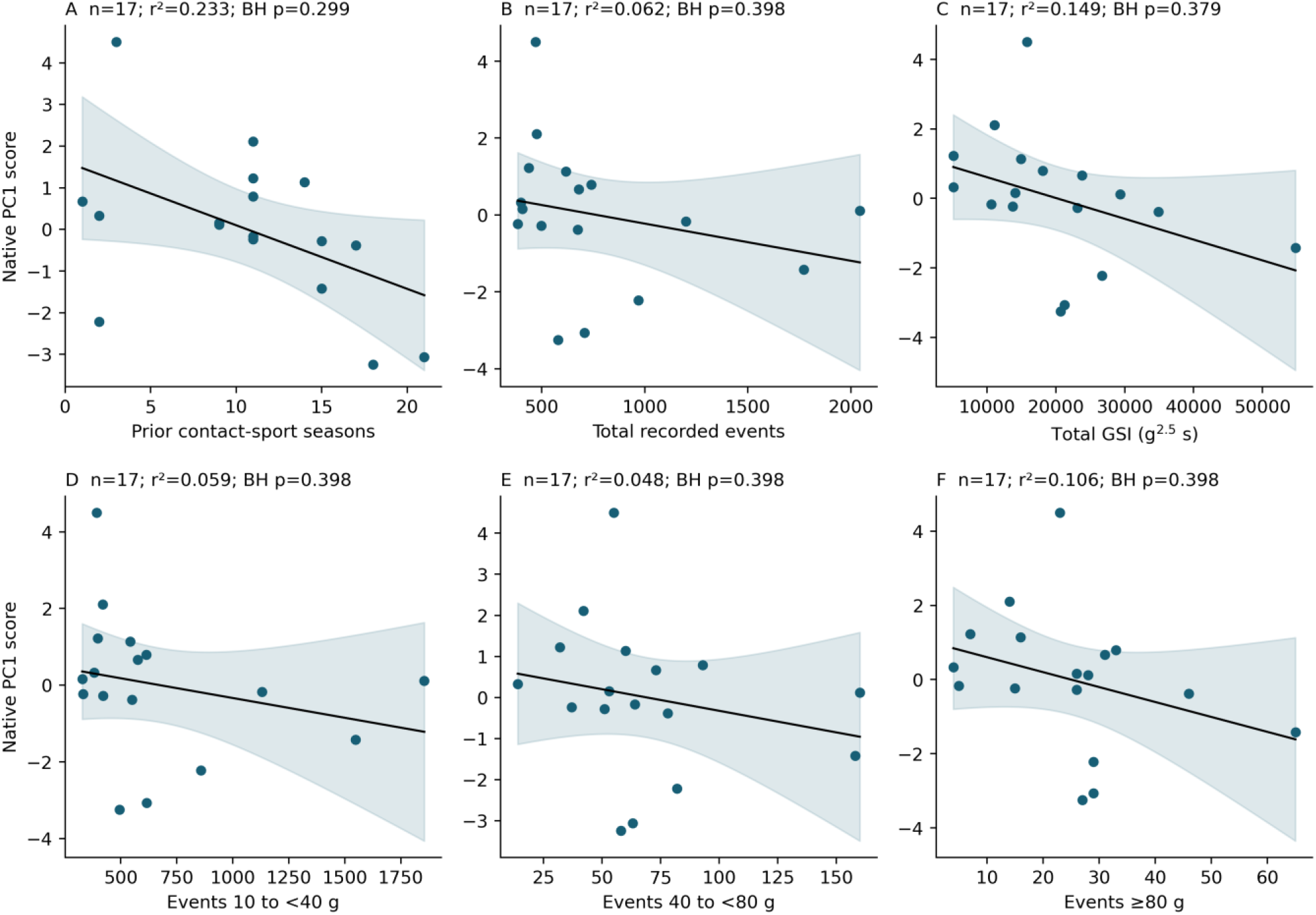
Six exposure associations with native PC1, n=17. Lines are ordinary least-squares fits and shaded bands show marginal 95% confidence intervals for the mean response. PC1 represents mixed changes across cognition, symptoms, and balance. None of six associations survived BH adjustment.

### Exploratory prediction results

In exploratory nested cross-validation, the six exposure measures did not provide useful out-of-sample prediction of the fixed cognitive-change component (median repeated r=−0.023; predictive R^2^=−0.204 against the overall-mean benchmark and −0.050 against the outer-training-mean benchmark). Sensitivity analyses produced the same overall conclusion (Supplementary Table S5).

## Discussion

Recorded helmet-sensor exposure and repeated cognitive performance were characterised in British amateur American football. Several individual tests improved, but none of six PC1 exposure associations survived BH adjustment. Prior seasons had the strongest nominal association among the tested exposures, with substantial uncertainty and mixed component interpretation. Its sign does not support the interpretation that greater prior exposure was associated with worse global cognition based on this study.

Improvement is compatible with familiarisation and other influences on repeated assessment [32,33]; these data do not identify the contribution of practice. PC1 describes between-participant differences in a centred pattern of change. Relatively less favourable change need not mean absolute deterioration, and here the mixed loadings oppose some favourable changes to others. Treating PC1 as a uniform global worsening score would overstate the measured construct.

Prior seasons provide a pragmatic history measure that also reflects experience and selection [34]. This cohort does not establish an adjusted association with an interpretable global cognitive outcome.

Uncertain exposure associations and the absence of useful prediction in this small exploratory cohort do not establish absence of neurological effects or safety. Imaging and physiological endpoints examine different aspects of brain function [16–18,35,36]. Outcome definition, measurement quality, and recording completeness may contribute to divergent findings. Scoring and component sensitivities are reported transparently rather than selected by statistical significance.

Interpretation is limited by the small, linked cohort and uncertain exposure completeness. Devices measured linear acceleration only, and helmet coupling differs from head centre-of-mass motion [37,38]. Broad session windows do not remove event-level artefacts. Participant activation and missed recordings can underestimate exposure; non-impact recordings can inflate counts or GSI. Prior seasons also reflect experience, selection, and playing history.

There was no non-contact comparison group to quantify familiarisation. Task programme versions were incompletely documented, and task presentation timings were not recorded. All participants were adult males, limiting generalisability. Non-significant results do not establish absence of an effect, injury, or harm.

Future work should use larger, prospectively specified cohorts, verified playing-session records, and measurements incorporating linear and rotational kinematics. The present study documents season-long monitoring in British amateur American football, with conclusions bounded by its recording and measurement limitations.

## Acknowledgements

We thank the participating players, teams, and University of Exeter research support staff.

## Data availability

Pseudonymised data supporting the findings and reproducible code are available from the corresponding author on reasonable request, subject to consent, ethics, and data-governance requirements. Identifiable participant information is not shared.

## Author contributions

Paul Weightman Potter and Bert Bond conceptualised the study, developed the methodology, and prepared the manuscript draft. Paul Weightman Potter provided study resources, prepared the visualisations, supervised and administered the project, and acquired funding. Paul Weightman Potter and Emily Rose validated the work and conducted the formal analyses. Paul Weightman Potter, Emily Rose, and Nicholas Tolliday conducted the investigation and collected data. Paul Weightman Potter, Emily Rose, and Nicholas Tolliday curated the data. All authors reviewed and edited the manuscript.

