## Supplementary materials for "Season-long head acceleration exposure, prior contact-sport history, and cognitive performance in British amateur American football players"

### Participant flow and scoring

Thirty players were recruited. Nine withdrew and were excluded entirely, and one had exposure data only. Twenty formed the cognitive cohort; 17 had linked exposure records. Reported exposure summaries describe these 17 linked participants. Mid-season observations were available for 17. Missing measurements were retained separately from withdrawal and sensor exclusions. An assessment conducted outside this study was excluded from the study visits.

The primary task-scoring approach was accompanied by a correct-response reaction time (RT) sensitivity analysis. Stroop used mean incongruent minus congruent RT without an accuracy or block-label filter; the sensitivity used correct responses. Go/No-Go errors were summed, and the primary RT definition excluded exactly 2,000 ms; the sensitivity used correct Go responses only. Every 2-back file contained 75 trials; errors were calculated as 75 minus summed correctness, and RT was calculated from correct trials, excluding those of exactly 3,000 ms. Visual search used correct target-present trials identified by the prespecified task flag within each array size. No additional trimming, timeout rule, or practice exclusion was introduced.

**Table S1. Participant flow and analysis sets.**

| Stage or set | n |
|---|---|
| Recruited | 30 |
| Withdrawn; excluded entirely | 9 |
| Exposure data only; not included in reported summaries | 1 |
| Seasonal cognitive cohort | 20 |
| Mid-season cognitive visits | 17 |
| Linked exposure/cognition cohort | 17 |

**Table S2. Primary-scoring seasonal omnibus tests. REML random-intercept models, Satterthwaite F tests, and BH adjustment across 17 individual outcomes.**

| Outcome | F | df num. | df denom. | Raw p | BH p |
|---|---|---|---|---|---|
| mBESS errors | 0.38 | 2 | 35.43 | 0.6888 | 0.6888 |
| Concentration | 3.69 | 2 | 35.67 | 0.0348 | 0.0592 |
| Delayed recall | 7.99 | 2 | 35.27 | 0.0014 | 0.0078 |
| Digits backward | 3.31 | 2 | 35.70 | 0.0479 | 0.0741 |
| Go/No-Go errors | 0.59 | 2 | 36.61 | 0.5615 | 0.5966 |
| Go/No-Go RT (ms) | 1.68 | 2 | 36.19 | 0.2013 | 0.2632 |
| Immediate memory | 6.33 | 2 | 35.91 | 0.0044 | 0.0157 |
| 2-back errors | 5.67 | 2 | 35.51 | 0.0073 | 0.0177 |
| 2-back RT (ms) | 6.34 | 2 | 33.41 | 0.0046 | 0.0157 |
| Orientation | 5.83 | 2 | 42.17 | 0.0058 | 0.0165 |
| Search RT 10 items (ms) | 4.94 | 2 | 35.64 | 0.0128 | 0.0272 |
| Search RT 15 items (ms) | 19.32 | 2 | 35.39 | 2.1e-06 | 3.6e-05 |
| Search RT 20 items (ms) | 16.78 | 2 | 35.53 | 7.4e-06 | 6.3e-05 |
| Search RT 5 items (ms) | 3.84 | 2 | 35.88 | 0.0308 | 0.0582 |
| Stroop interference (ms) | 0.69 | 2 | 35.93 | 0.5063 | 0.5739 |
| Symptom count | 2.47 | 2 | 35.62 | 0.0991 | 0.1404 |
| Symptom severity | 1.51 | 2 | 35.76 | 0.2348 | 0.2851 |

Orientation had near-boundary participant variance. Baseline and post-season scores were constant at 5, so the omnibus effect concerns mid-season departures. Model contrasts use the estimated pooled covariance.

**Table S3. Baseline-to-post-season model contrasts. Changes are post-season minus baseline in original outcome units. Confidence intervals are marginal 95% intervals. P values use Holm–Šídák across three time contrasts within each outcome.**

| Outcome | Post minus baseline | 95% CI | Adjusted p |
|---|---|---|---|
| mBESS errors | 0.500 | -1.056 to 2.056 | 0.8065 |
| Concentration | 0.500 | 0.035 to 0.965 | 0.0701 |
| Delayed recall | 1.150 | 0.566 to 1.734 | 0.0009 |
| Digits backward | 0.500 | 0.038 to 0.962 | 0.0956 |
| Go/No-Go errors | 0.150 | -0.281 to 0.581 | 0.7344 |
| Go/No-Go RT (ms) | -34.423 | -72.736 to 3.889 | 0.213 |
| Immediate memory | 2.350 | 0.844 to 3.856 | 0.0095 |
| 2-back errors | -3.100 | -5.130 to -1.070 | 0.0113 |
| 2-back RT (ms) | -115.262 | -183.050 to -47.475 | 0.0045 |
| Orientation | 0.000 | -0.152 to 0.152 | 1 |
| Search RT 10 items (ms) | -99.469 | -165.759 to -33.180 | 0.0131 |
| Search RT 15 items (ms) | -182.149 | -249.057 to -115.241 | 9.6e-06 |
| Search RT 20 items (ms) | -272.105 | -371.895 to -172.315 | 9.4e-06 |
| Search RT 5 items (ms) | -98.612 | -171.041 to -26.183 | 0.0268 |
| Stroop interference (ms) | 15.012 | -35.980 to 66.004 | 0.7923 |
| Symptom count | -1.250 | -2.518 to 0.018 | 0.1511 |
| Symptom severity | -1.950 | -4.529 to 0.629 | 0.3503 |

**Table S3b. Correct-response RT sensitivity: changed outcomes only, with the same contrast definitions.**

| Outcome | Post minus baseline | 95% CI | Adjusted p |
|---|---|---|---|
| Go/No-Go RT (ms) | -34.172 | -72.184 to 3.839 | 0.2125 |
| Stroop interference (ms) | 13.920 | -37.717 to 65.557 | 0.7079 |

**Table S4. Exposure regressions using native PC1, n=17. Each scoring variant has a separate six-test BH family. Confidence intervals are marginal 95% intervals.**

##### Primary scoring

| Exposure | Slope (95% CI) | $r^2$ | Raw p | BH p |
|---|---|---|---|---|
| Prior seasons | -0.15274 (-0.30529 to -0.00018) | 0.233 | 0.0498 | 0.2986 |
| Total events | -0.00097 (-0.00304 to 0.00110) | 0.062 | 0.3347 | 0.3984 |
| Total GSI | -6e-05 (-0.00014 to 1.9e-05) | 0.149 | 0.1264 | 0.3792 |
| Events 10 to <40 g | -0.00103 (-0.00330 to 0.00124) | 0.059 | 0.3484 | 0.3984 |
| Events 40 to <80 g | -0.01052 (-0.03631 to 0.01527) | 0.048 | 0.3984 | 0.3984 |
| Events ≥80 g | -0.04028 (-0.10458 to 0.02402) | 0.106 | 0.2017 | 0.3984 |

##### Correct-response RT sensitivity

| Exposure | Slope (95% CI) | $r^2$ | Raw p | BH p |
|---|---|---|---|---|
| Prior seasons | -0.16147 (-0.30916 to -0.01378) | 0.266 | 0.0342 | 0.205 |
| Total events | -0.00100 (-0.00304 to 0.00105) | 0.067 | 0.3152 | 0.3407 |
| Total GSI | -6.2e-05 (-0.00014 to 1.6e-05) | 0.162 | 0.1096 | 0.3115 |
| Events 10 to <40 g | -0.00105 (-0.00329 to 0.00119) | 0.062 | 0.3337 | 0.3407 |
| Events 40 to <80 g | -0.01170 (-0.03705 to 0.01364) | 0.061 | 0.3407 | 0.3407 |
| Events ≥80 g | -0.04403 (-0.10682 to 0.01876) | 0.130 | 0.1558 | 0.3115 |

### Exploratory prediction

These exploratory penalised linear regression models, fitted using glmnet, did not demonstrate useful out-of-sample prediction; the details below are provided for transparency. Outer evaluation used five folds repeated 20 times and inner tuning four folds repeated five times. Seed: 20261023. The mixing parameter alpha was 0, 0.25, 0.5, 0.75, or 1, spanning ridge, elastic-net, and lasso regression. The penalty parameter lambda comprised 60 logarithmically spaced values from 0.0001 to 100. Predictor transformations and training-target scaling were fitted inside each split. Hyperparameters were selected by minimising validation mean squared error (MSE); ties were resolved by preferring larger lambda, followed by larger alpha. Repeats are dependent resampling summaries, not confidence intervals.

The fixed-PC1 evaluation used a full-cohort target, including the mean-imputed 2-back RT change, and served as a conditional benchmark with 17 evaluable participants. The fixed-PC1 models used all six exposure predictors. The training-derived PCA sensitivity omitted total events because it equals the sum of the three event-category counts, leaving five predictors. In that sensitivity, incomplete held-out outcome profiles were not imputed into observed targets, leaving 16 evaluable participants. Training-derived PCs retained their native signs without a common polarity, so pooled r, squared correlation, and overall-mean $R^2$ were unavailable. Procedural errors are expressed in training-PC SD units; fixed-target errors are expressed in raw full-cohort PC1 units. These errors cannot be directly compared.

**Table S5. Held-out performance. Correlation and $R^2$ columns are medians of repeat metrics. RMSE is the square root of the mean repeat MSE; MAE is the mean repeat MAE. Overall and training refer to the reference predictions used for the $R^2$ denominator.**

| Target/scoring | n | r | $r^2$ | $R^2$ overall | $R^2$ training | RMSE | MAE |
|---|---|---|---|---|---|---|---|
| Correct-response RT; fixed | 17 | 0.043 | 0.005 | -0.176 | -0.027 | 2.051 | 1.474 |
| Correct-response RT; training PCA | 16 | — | — | — | 0.007 | 0.609 | 0.437 |
| Primary; fixed | 17 | -0.023 | 0.007 | -0.204 | -0.050 | 2.072 | 1.524 |
| Primary; training PCA | 16 | — | — | — | 0.001 | 0.623 | 0.471 |

For each repeat, $r=\mathrm{cor}(y,p)$, squared correlation$=r^2$, predictive $R^2=1-\Sigma(y-p)^2/\Sigma(y-\mathrm{mean}(y))^2$, training-baseline $R^2=1-\Sigma(y-p)^2/\Sigma(y-\text{training mean})^2$, RMSE was the square root of $\mathrm{mean}((y-p)^2)$, and MAE was $\mathrm{mean}(\mathrm{abs}(y-p))$. Median $r^2$ need not equal squared median r.

### PCA interpretation and sensitivity

Full-cohort PCA used 17 linked players, 16 varying changes, and one mean-imputed 2-back RT change; the resulting PC1 scores were used in exposure regressions and the fixed-target prediction benchmark. Inputs were oriented once so favourable change was positive; native PC1 was not reversed. Mixed weights prevent a uniform better/worse interpretation. Digits and concentration overlapped. The domain-balanced sensitivity analysis used eight domains of centred change, including symptoms and balance. The domain-balanced sensitivity score was calculated from the same oriented, standardised changes used for PCA. Missing standardised values were replaced with zero, corresponding to the observed cohort mean change. Changes were averaged within each of eight domains: symptoms, memory, concentration, balance, Stroop, Go/No-Go, working memory, and visual search. The eight domain means were then averaged with equal weight. Digits backward was excluded as a separate measure to avoid overlap with concentration; orientation was excluded from this sensitivity score. This score measures relative change within the cohort, with higher values indicating more favourable relative change, and is not a measure of absolute cognitive improvement. Six exposure regressions were fitted for each scoring variant, with BH adjustment within each six-test family.

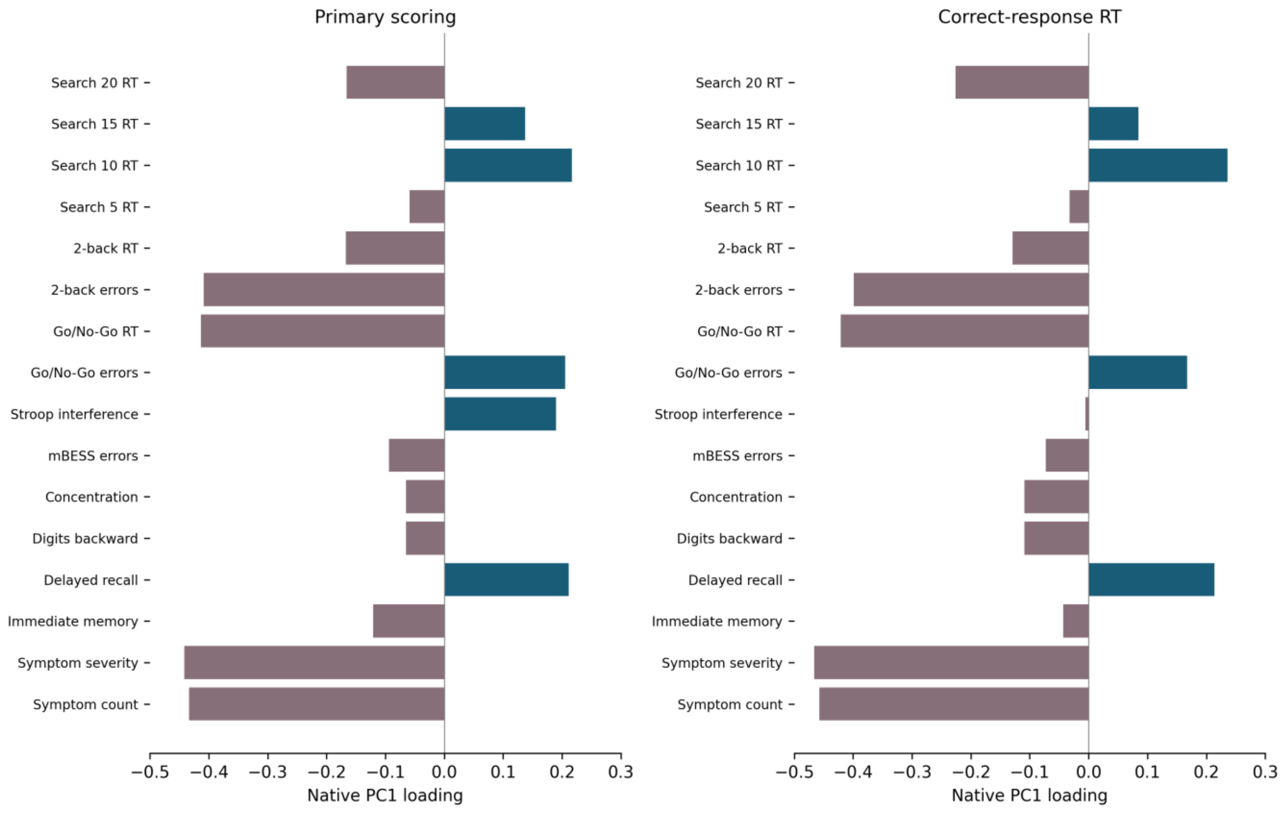


**Figure S1. Native PC1 loadings for primary scoring and the correct-response RT sensitivity analysis. Signed loadings show the mixed change pattern.**

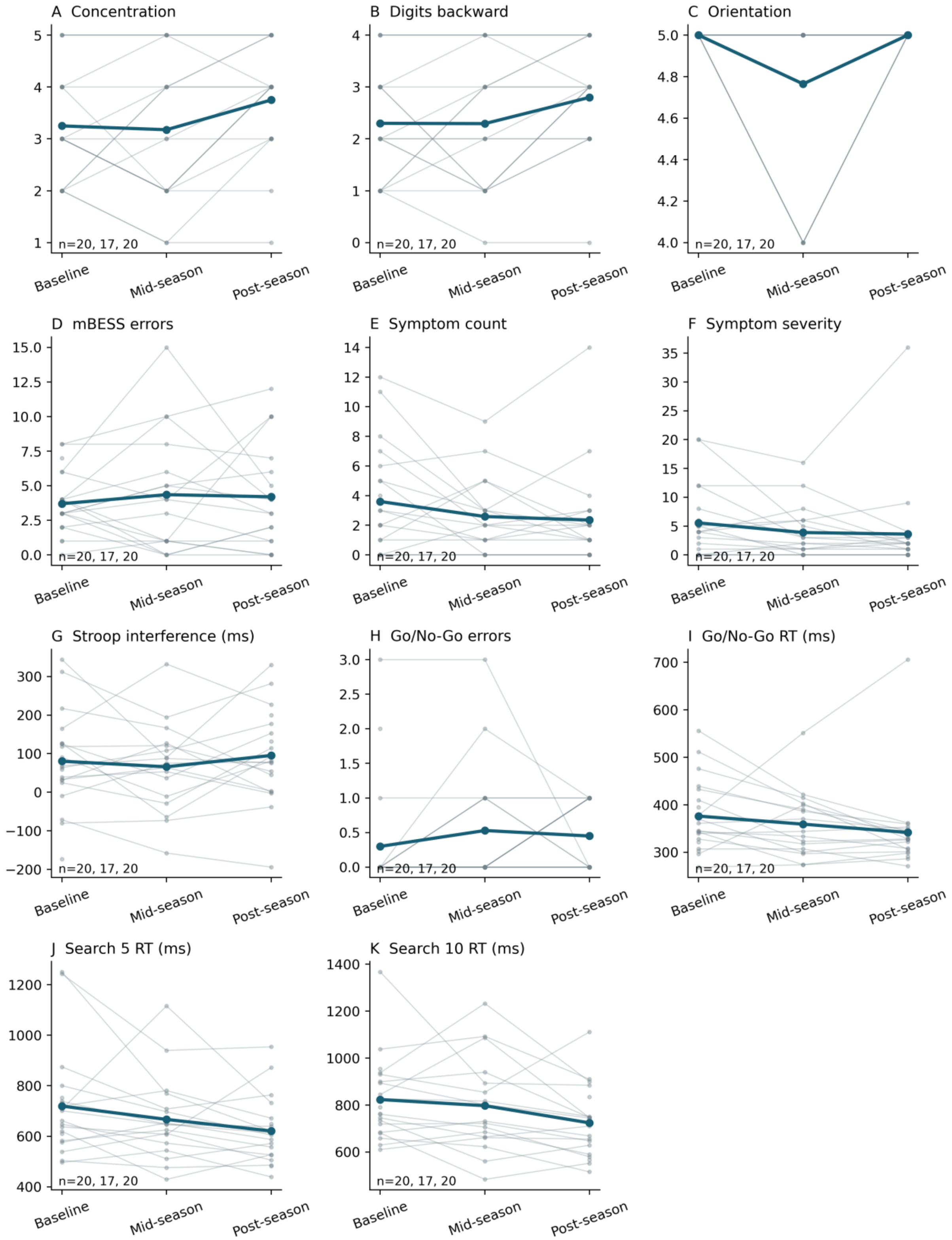


**Figure S2. Additional individual outcomes. Points and thin lines show participants; thick lines show observed means. Sample sizes are outcome-specific. These plots are descriptive; inferential analyses and multiplicity adjustments are reported separately in Tables S2 and S3.**

#### Table S6. Domain-balanced sensitivity analyses

Each variant contains six separate simple linear regressions (n=17). Slopes are in domain-score units per unit of exposure; confidence intervals are marginal 95% intervals. BH adjustment is applied separately within each scoring variant. No association survives adjustment.

##### Primary scoring

| Exposure | Slope (95% CI) | $r^2$ | Raw p | BH p |
|---|---|---|---|---|
| Prior seasons | 0.02029 (-0.001115 to 0.0417) | 0.214 | 0.0616 | 0.3694 |
| Total events | 4.21e-05 (-0.0002531 to 0.0003373) | 0.006 | 0.7653 | 0.8150 |
| Total GSI | 6.016e-06 (-5.361e-06 to 1.739e-05) | 0.078 | 0.2774 | 0.5549 |
| Events 10 to <40 g | 3.616e-05 (-0.0002876 to 0.0003599) | 0.004 | 0.8150 | 0.8150 |
| Events 40 to <80 g | 0.0009093 (-0.00272 to 0.004539) | 0.019 | 0.6012 | 0.8150 |
| Events ≥80 g | 0.006117 (-0.00269 to 0.01492) | 0.127 | 0.1594 | 0.4783 |

##### Correct-response RT sensitivity

| Exposure | Slope (95% CI) | $r^2$ | Raw p | BH p |
|---|---|---|---|---|
| Prior seasons | 0.02229 (0.000754 to 0.04383) | 0.245 | 0.0434 | 0.2603 |
| Total events | 0.0001081 (-0.00019 to 0.0004062) | 0.038 | 0.4516 | 0.4855 |
| Total GSI | 6.8e-06 (-4.772e-06 to 1.837e-05) | 0.095 | 0.2296 | 0.4591 |
| Events 10 to <40 g | 0.0001099 (-0.0002176 to 0.0004373) | 0.033 | 0.4855 | 0.4855 |
| Events 40 to <80 g | 0.001584 (-0.002074 to 0.005243) | 0.054 | 0.3706 | 0.4855 |
| Events ≥80 g | 0.006285 (-0.002755 to 0.01532) | 0.128 | 0.1591 | 0.4591 |
